# Proton selective conductance and gating of lysosomal cation channel TMEM175

**DOI:** 10.1101/2024.11.01.621492

**Authors:** Tobias Schulze, Timon Sprave, Carolin Groebe, Jan Krumbach, Magnus Behringer, Kay Hamacher, Gerhard Thiel, Christian Grimm, Oliver Rauh

## Abstract

The lysosomal cation channel TMEM175 is crucial for maintaining lysosomal function and pH homeostasis, and its aberrant function is linked to Parkinson’s disease (PD). While TMEM175 activity was first interpreted in the context of its potassium (K^+^) selective conductance, subsequent studies revealed also a substantial permeability to protons (H^+^). Here we dissect the complex changes in TMEM175 conductance and current reversal voltages in response to pH jumps on the luminal side of the channel protein. In whole-cell patch clamp experiments with plasma membrane redistributed TMEM175 we show that a pH jump from symmetrical pH 7.4 to pH 4.7 on the luminal side triggers a continuous rise in inward and outward current, concomitant with a transient positive excursion of the reversal voltage (E_rev_). The peak E_rev_ shift remains almost 100 mV below the estimated equilibrium voltage for protons and shows little sensitivity to the K^+^ gradient. The data are consistent with a scenario in which a TMEM175 mediated proton flux elicits a fast collapse of the pH gradient. In MD simulations we identify the luminal H57 as titratable partner for the formation of intra- and inter-subunit salt bridges with D279 and E282 for stabilizing the channel open state. This presumed gating function is confirmed by mutational studies and lysosomal patch-clamp experiments in which a H57Y mutant exhibits a reduced pH dependency of activation. Our findings contribute to a better comprehension of TMEM175’s complex electrophysiological properties and foster understanding of TMEM175 as a pharmacological target for neurodegenerative disease therapy.

## Introduction

The transmembrane protein 175 (TMEM175) is ubiquitously expressed in the membrane of endosomes and lysosomes. Accumulating evidence suggests that its dysfunction is a risk factor for the development of Parkinson’s disease (PD). For example, TMEM175 missense mutations like M393T and Q65P, which are frequently found in PD patients, impact the risk for PD in humans. This may be causally linked to corrupted key lysosomal functions which are observed in experimental knockout studies of the TMEM175 protein. The consequent impaired degradation of lysosomal content such as α-synuclein oligomers can be causally related to an augmented risk for PD (Xie & Hu, 2022). While the physiological consequences of TMEM175 dysfunction are becoming increasingly evident, the primary function of this transmembrane protein is still a matter of debate. In search for an ion channel, which mediates a postulated K^+^ conductance over lysosomal membranes, TMEM175 was initially described as a lysosomal K^+^ channel (Cang et al., 2015). In agreement with the view that TMEM175 provides the main K^+^ conductance across the lysosomal membrane, the lysosomal membrane loses its K^+^ conductance and depolarizes upon pharmacological inhibition or genetic knockout of TMEM175. The finding that this loss of K^+^ conductance leads, under different physiological conditions to hyper-acidification or alkalinization of lysosomes, suggested that the K^+^ conductance is important for maintaining a stable lysosomal pH (Cang et al., 2015; Jinn et al., 2017). Subsequent studies were able to resolve the atomistic structure of the human TMEM175 channel and its bacterial homologs (Brunner et al., 2020; C. Lee et al., 2017; Oh et al., 2022, 2024). These studies revealed a conserved architecture in which a central pore is formed in the bacterial channel by an assembly of homotetramers. Each protomer is composed of six transmembrane domains (TMDs) of which the first two N-terminal TMDs contribute to the formation of the pore. In the mammalian protein, the same overall channel architecture is achieved by an assembly of homodimers in which each of the monomers resembles a duplication of the 6TMD architecture of prokaryotic proteins. When bacterial or human TMEM175 proteins were isolated in the presence of KCl it was possible to identify distinct non-protein densities in a central ion-conduction pore. In agreement with a K^+^ channel function, these densities could be assigned to K^+^ ions (Brunner et al., 2020). In follow-up studies, the conductive properties of hTMEM175 were further investigated in solutions with an acidic pH on the luminal side (pH_L_) i.e., the side facing the acidic milieu of the lysosomal lumen. Several studies have confirmed that the TMEM175 conductance is dominated by a K^+^ conductance at neutral pH. However, an acidic pH_L_ causes not only a strong increase in TMEM175 conductance but also a shift in ion selectivity towards a preference for H^+^ ions (Hu et al., 2022; Zheng et al., 2022). Support for a scenario in which the TMEM175 conductance is dominated by a H^+^ current at acidic pH_L_ comes from fluorometric measurements showing a strong acidification of the lysosomal pH as a consequence of channel activation (Bazzone et al., 2023; Hu et al., 2022; Zhang et al., 2023). Complementary electrophysiological studies of TMEM175 currents furthermore demonstrated that the reversal voltage (E_rev_) of this channel shifts in patch-clamp recordings with changes of the transmembrane pH gradient (ΔpH). While the direction of the ΔpH evoked shift in E_rev_ is in agreement with a selective H^+^ conductance, the reported absolute E_rev_ voltages are in most cases well below the expected 60 mV per pH unit. Only in one study, the E_rev_ follows, at very low pH values the theoretically expected shift of E_H+_ (Zheng et al., 2022). In that case, however, the extrapolated E_rev_ value is well negative of 0 mV for symmetrical pH buffers e.g., the expected value in the absence of a driving force. A strict biophysical interpretation of these data suggests that the channel may not only transport H^+^ but also other ions. An alternative interpretation is that the measured reversal voltage is not identical with the calculated Nernst voltage for H^+^ due to experimental challenges. The latter interpretation would be reasonable considering the known problem of determining the correct Nernst voltage for H^+^ in cells. It has been shown in mammalian cells expressing the proton selective M2 channel, that rapid acidification of the external buffer results in a short-lasting positive shift of the reversal voltage towards the expected H^+^ equilibrium voltage before the reversal voltage gradually moves within seconds back to more negative values (Mould et al., 2000). A systematic investigation of this phenomenon has shown that the influx of H^+^ into cells generates a rapid and long-lived accumulation of H^+^ ions at the membrane/cytosol interface. The degree and the velocity of this dynamic change in H^+^ reversal voltage is augmented and accelerated by an increasing driving force for the H^+^ inward current (Mould et al., 2000). Hence, in patch-clamp recordings the prevailing H^+^ equilibrium voltage is no longer defined by the given pH of the external buffer and the patch pipette. Even high concentrations of H^+^ buffers in the pipette solution are not able to overcome this problem in conventional patch-clamp experiments (Mould et al., 2000). The same study shows that such dynamic variation of the H^+^ reversal voltage in cells is not only a property of the M2 protein but is also generated by the H^+^ conducting uncoupler FCCP. This implies that the H^+^ reversal voltage will change over time in response to any H^+^ conductance including the TMEM175 channel. In the published studies on the H^+^ conductance of TMEM175 this phenomenon of a dynamic reversal voltage has not yet been reported (Hu et al., 2022; Zhang et al., 2023; Zheng et al., 2022). Hence, the physicochemical meaning of the published reversal voltages of a TMEM175 generated conductance in an experimental system with inevitable dynamic changes in the H^+^ reversal voltage is not clear. Yang et al. (2023) have recently reported that a pH jump from 7.2 to 4.6 and back to 7.2 causes transient TMEM175 currents in whole-endolysosomal patch-clamp recordings (Yang et al., 2023). The latter were recorded under voltage clamp conditions at one given voltage and interpreted as evidence for a scenario in which a H^+^ current into the lysosomal lumen causes an acidification, which eventually reduces TMEM175 activity via pH dependent gating. Without any experimental evidence for such a pH dependent gating the transient currents can also be interpreted simply in the context of dynamic changes in the driving force for H^+^ currents. A pH jump to an acidic buffer can elicit an initial increase in the H^+^ current, which then decays with a progressive acidification of the lysosomal lumen. Such a simple thermodynamic interpretation of the data is supported by the experimental finding that the kinetics of the ΔpH induced transient currents depend on the clamp voltages (Yang et al., 2023).

Inspired by the work on the M2 channel we evaluated here the relationship between the H^+^ conductance of the TMEM175 protein and changes in the reversal voltage. By altering the ionic composition of the pipette and buffer solutions, our experiments revealed that acidification of the buffer on the luminal side of the channel protein induces a transient positive shift in E_rev_, indicating a predominant H^+^ conductance followed by a shift in E_rev_ back to values close to 0 mV. Additional experiments could show that this backshift is linked to a decreasing proton gradient in the microenvironment close to the membrane. Importantly, mutational studies based on TMEM175’s open and closed conformation structures revealed one amino acid, H57 to function as pH sensor in TMEM175, which was confirmed by lysosomal patch-clamp electrophysiology. We thus provide new evidence for the dual conductance of TMEM175, supporting a critical role of TMEM175 for lysosomal acidification and by identifying a key amino acid acting as the pH sensor in TMEM175. Our data will thus have important implications for our understanding of channel gating by protons and the development of small molecule drugs for the treatment of neurodegenerative diseases such as PD.

## Results and Discussion

To analyze the time course of pH induced conductance changes in TMEM175, the protein was transiently overexpressed in HEK293 cells. This causes insertion of the functional TMEM175 channel into the plasma membrane (Brunner et al., 2020) with an orientation in which the side facing the lumen of the lysosome is exposed to the extracellular solution. In this orientation TMEM175 currents can be easily monitored in the whole cell configuration before, during and after the exchange of the extracellular buffer (= luminal solution). Empty vector transfected HEK293 cells served as a negative control.

### Acidification of the luminal side induces transient shift of reversal voltage to positive values

Fig. 1A shows an exemplary recording from a control cell, which was repetitively challenged every 10 seconds by voltage ramps (± 120 mV; 0.46 mV/ms) in symmetrical buffers with 140 mM potassium methanesulfonate (KCH_3_SO_3_; K-MS) and pH 7.4. Exemplary current responses to selected voltage ramps (Fig. 1A) as well as current densities, membrane conductance and the current reversal voltages (E_rev_) (Fig. 1C) from all ramps reveal a low conductance in pH 7.4 and an E_rev_ value close to 0 mV. A rapid exchange of the external pH from pH 7.4 to pH 4.7 has, over a recording window of 3 min no perceivable impact neither on conductance nor on E_rev_. This overall insensitivity of both parameters to a pH jump was confirmed in 12 other experiments of the same kind. The individual time-courses as well as the mean values are shown for E_rev_ in Fig. 1E.

**Fig. 1.**
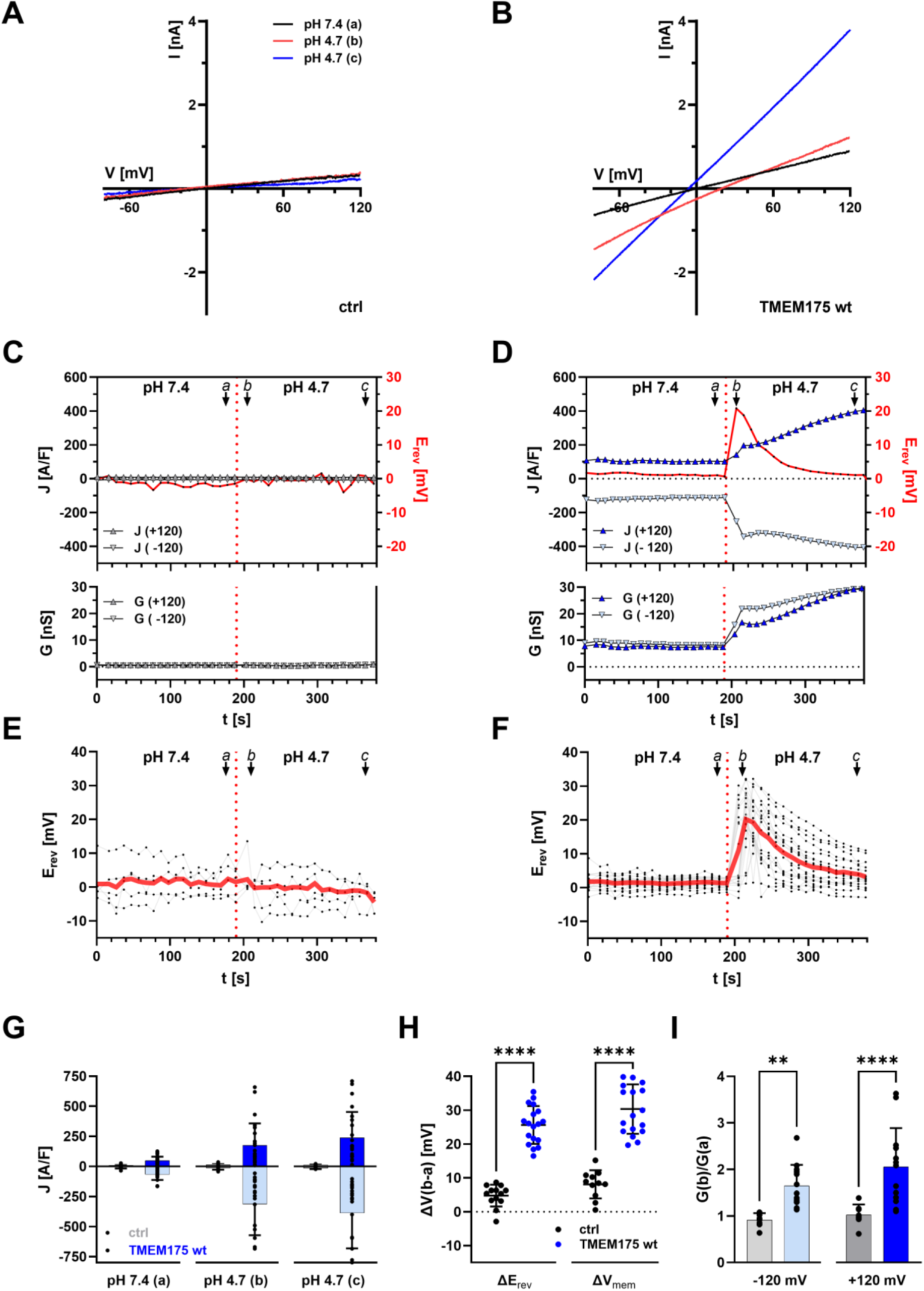
Acidification of the luminal side induces a continuous increase in conductance but only a transient shift of E_rev_ to positive values. **(A,B)** Representative current responses to voltage ramps from +120 mV to −120 mV of empty vector transfected (A) and TMEM175 expressing (B) HEK293 cells recorded shortly before (a, black), shortly after (b, red) and 3 min after (c, blue) a pH jump from 4.7 to 7.4 in the bath solution (luminal side). Bath and pipette solution contained 140 mM K-MS. The internal pH was 7.4. The external pH is indicated at the top. **(C,D)** Representative time-courses of reversal voltage (E_rev_) as well as current densities (upper graph) and chord conductance (lower graph) at +/-120 mV of empty vector transfected (C) and TMEM175 expressing (D) HEK293 cells. Values were taken from voltage-ramp recordings as in (A) and (B). **(E,F)** Individual and averaged time-courses of E_rev_ of empty vector transfected (E) and TMEM175 expressing (F) HEK293 cells from pH jump experiments as in (C) and (D). **(G)** Current densities at +/-120 mV at time points a, b and c in (C-F). Bars represent arithmetic mean ± SD. Values from individual recordings are shown as black closed circles. **(H)** Maximal change in reversal voltage E_rev_ and free running membrane voltage V_mem_ in response to pH jump from 7.4 to 4.7 of empty vector transfected (black) and TMEM175 expressing (blue) HEK293 cells. Arithmetic mean is shown as horizontal bar. Values from individual recordings are shown as closed circles. **(I)** pH-jump induced change in ion conductance at -/+120 mV for empty vector transfected (gray) and TMEM175 expressing (blue) HEK293 cells. Bars represent arithmetic mean ± SD. Values from individual recordings are shown as black closed circles. Statistical comparisons in (H) and (I) were made using an unpaired two-tailed Student’s t-test (**: p < 0.01, ***: p < 0.001, ****: p < 0.0001).

The same procedure was repeated with cells expressing TMEM175. At pH 7.4 the E_rev_ value was also close to 0 mV; the conductance however was ca. 3.3 fold elevated over that in control cells. Rapid acidification of the external solution from pH 7.4 to pH 4.7 elicited a progressive increase in conductance and caused a rapid shift of the reversal voltage. The exemplary ramp currents (Fig. 1B) as well as the continuous recording of the current densities and membrane conductance (Fig. 1D) show that both progressively increases over time. A scrutiny of the reversal voltages of the individual ramps as well as the continuous monitoring of this value shows that the pH jump elicits an excursion of this value from 0 mV to a peak at ∼+20 mV before the value decays back towards 0 mV (Fig. 1D). The same response pattern with a transient shift of E_rev_ to positive voltages way below the calculated E_H_^+^ value and a drastic increase in inward and outward currents has been recorded in several (n = 25) analogous experiments (Fig. 1F&G).

One reason for the dynamic shift of the reversal voltage could be related to the constant application of voltage ramps. Frequent challenging of cells with high voltages might favor a flux of ions and a consequent change in the local ionic gradient for the transported ions. To minimize the directional effect of the applied voltage ramps on the pH gradient, the ramps were applied in both directions (Supplementary Fig. S1). In fact, the forward ramp (from +120 mV to −120 mV) and the reverse ramp (from −120 mV to +120 mV) show different reversal potentials (Fig. S1). The reversal potential of the reverse ramp is shifted to the left by about 5 mV compared to the forward ramp (Supplementary Fig. S1B,C), which indicates that the inward H^+^ current indeed reduces the pH gradient. For this reason, all E_rev_ values reported below represent the average of the E_rev_ values of the forward and reverse ramps.

To further test the impact of frequent voltage ramps on the dynamics of E_rev_ we also measured in control and TMEM175 expressing cells the free running membrane voltage in the current clamp mode before and after acidifying the external pH from 7.4 to 4.7. The recordings show that this procedure elicits a similar transient depolarization in TMEM175 expressing cells (Supplementary Fig. S2). These results were confirmed in recordings in 18 other cells underscoring that the data are not depending on the recording method (Fig. 1H).

### Contribution of H^+^ and K^+^ fluxes to transient shift of reversal voltage

The transient nature in the shift of the E_R_ value in response to a jump from neutral to acidic pH has not been reported before from experiments under similar conditions with TMEM175 in the plasma membrane of HEK293 cells. In the following experiments, we attempted to find an explanation for this phenomenon. One possible cause of the transient excursion of E_rev_ towards E_H+_ (+160 mV) in TMEM175 expressing cells could be that the pH induced H^+^ conductance is only short lived. This explanation however can be excluded as the pH induced conductance of the membrane remains elevated or keeps even increasing after the E_rev_ value has reached its maximum (Fig. 1D&G).

An alternative explanation for the phenomenon is that a pH jump elicits a rapid H^+^ conductance followed by a delayed rise in K^+^ conductance. Such a sequential activation could, under the prevailing ionic conditions shift E_rev_ initially positive towards E_H_^+^ before eventually tending towards E_K_^+^ at 0 mV. To test if such a scenario was responsible for the transient excursion of E_rev_, we modified the experiments in Fig. 1; once E_rev_ had settled after the pH jump close to 0 mV, we reduced the external K^+^ concentration by a factor of 10. If the backshift of E_rev_ after the pH jump originates from a delayed and dominating K^+^ conductance, E_rev_ should settle in 14 mM K^+^ close to the theoretical K^+^ Nernst potential of −60 mV. Control experiments first confirmed that this maneuver had no appreciable effect on the membrane conductance of empty vector transfected HEK293 cells (Fig. 2C). But we still observed in the controls that E_rev_ shifted after the 10-fold drop in K^+^ on average ∼5 mV more negative (Fig. 2D). This indicates that endogenous K^+^ channels in HEK293 cells are not completely blocked by the canonical K^+^ channel blockers TEA^+^ and Cs^+^ in the external and pipette solution.

**Fig. 2.**
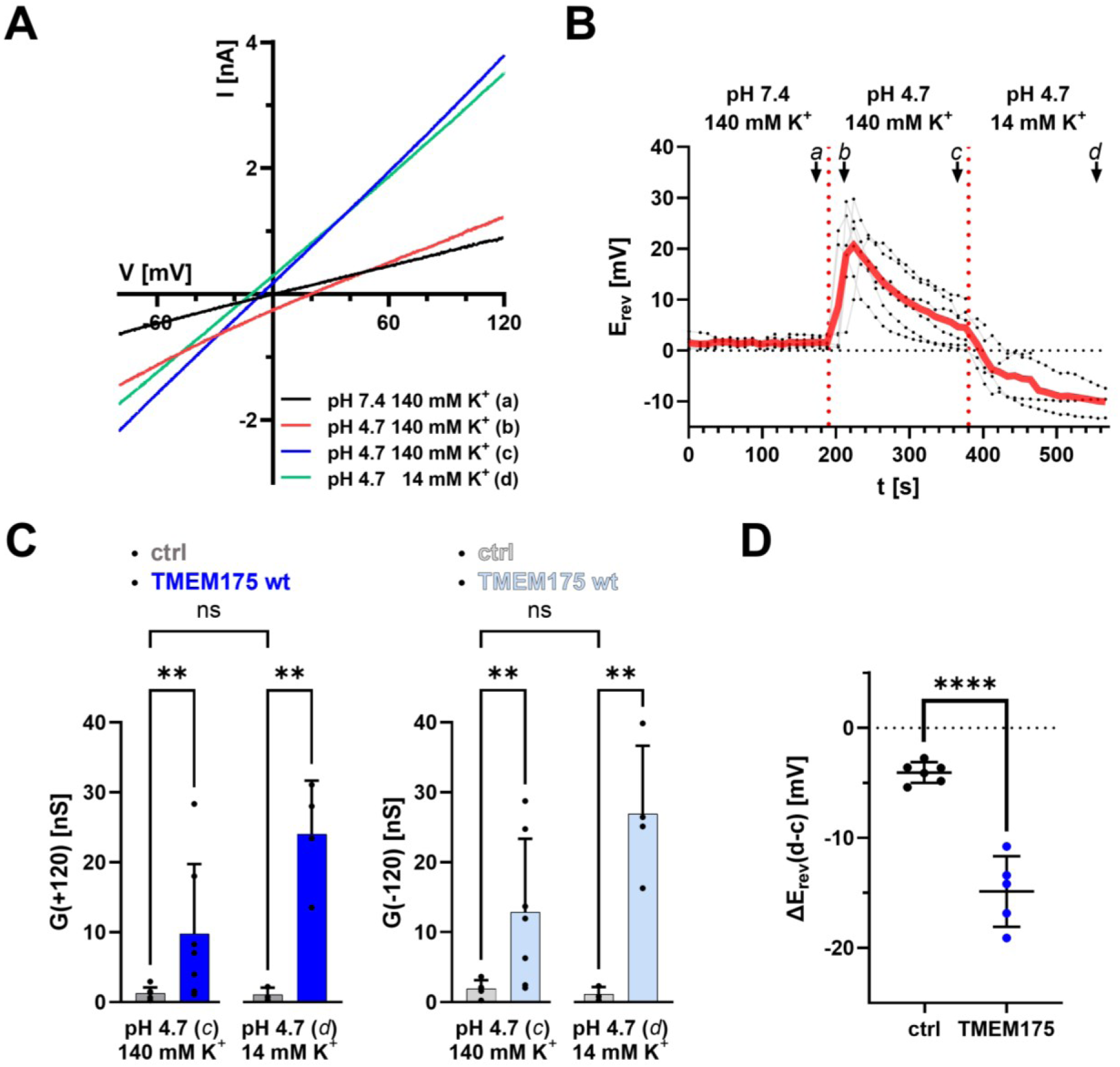
The steady-state conductance of TMEM175 after the pH jump is not dominated by K^+^. **(A)** Representative current responses to voltage ramps from +120 mV to −120 mV of TMEM175 expressing HEK293 cells recorded shortly before (a, black), shortly after (b, red) and 3 min after (c, blue) a pH jump from 4.7 to 7.4 in the bath solution (luminal side), and 3 min after reducing the K-MS concentration in the bath solution from 140 mM to 14 mM (d, geen). The internal pH was 7.4. **(B)** Individual and averaged time-courses of E_rev_ of TMEM175 expressing HEK293 cells. Values were taken from voltage-ramp recordings as in (A). The external pH is indicated at the top. **(C)** Chord conductances at +120 mV (left) and −120 mV (right) for empty vector transfected (gray) and TMEM175 expressing (blue) HEK293 cells at time points c and d in (B). Values were calculated from voltage-ramp recordings as in (A). Bars represent arithmetic mean ± SD. **(D)** Change in reversal voltage E_rev_ in response to a reduction of the external K^+^ concentration from 140 mM to 14 mM of empty vector transfected (black) and TMEM175 expressing (blue) HEK293 cells. E_rev_ values were calculated from voltage-ramp recordings as in (A) at time points c and d in (B). Statistical comparisons in (C) and (D) were made using an unpaired two-tailed Student’s t-test (**: p < 0.01, ****: p < 0.0001).

Fig. 2B shows that the same 10 fold drop in K^+^ shifted, in TMEM175 expressing cells E_rev_ not to the expected −60 mV; instead only a shift to about −15 mV was observed (Fig. 2B&D). These data clearly show that K^+^ currents are contributing but not dominating TMEM175 conductance in acidic buffer. Still the consequent prediction that E_rev_ must be dominated by the channel’s H^+^ conductance is only partially met by the experimental data. The measured E_rev_ increases transiently in the direction of E_H+_; but even the peak value remains in a buffer with pH 4.7 and symmetrical 140 mM K^+^ well below the theoretical expected H^+^ Nernst potential of +160 mV. We must therefore assume that the proton gradient rapidly collapses, like with other H^+^ conducting proteins (Mould et al., 2000), after TMEM175 activation. To test this hypothesis experiments as in Fig. 1 were repeated with a 10 fold increase in pH buffer concentration in pipette and bath solution (Supplementary Fig. S3). We expected that a higher buffer concentration might slow down the collapse of the pH gradient after TMEM175 activation. This should favor a stronger positive excursion of E_rev_ and a slower backshift towards 0 mV. At first glance and to our surprise, this was not the case. Instead, the pH-induced peak E_rev_ excursion was in the presence of 50 mM pH buffer with 15.9 ± 9.9 mV even smaller and the backshift towards 0 mV faster than in experiments with a low buffer concentration (25.6 ± 5.4 mV) (Supplementary Fig. S3C,E). In comparable experiments Mould and coworkers (Mould et al., 2000) have shown that even high buffer concentrations cannot adequately control, in patch clamp experiments the cytosolic pH. While this would explain an insensitivity of voltage changes to the pH buffer concentration it does not explain our finding that the pH induced E_rev_ excursion is even suppressed; for the moment we have no conclusive explanation for this unexpected result.

To further confirm that the backshift of E_rev_ towards 0 mV is solely caused by the collapse of the pH gradient in consequence of the proton conductivity of TMEM175, we repeated the experiments in Fig. 1 with solutions in which K^+^ was replaced by the large impermeable cation N-methyl-D-glucamine (NMDG^+^). In control cells a pH jump from pH 7.4 to 4.7 evoked no appreciable increase in current (Fig. 3C). The same experiments in TMEM175 expressing cells shows very low current densities in neutral buffer, which are, at positive and negative reference voltages (±120 mV) not appreciably different from control cells (Fig. 3C). After exchanging the pH from pH 7.4 to 4.7 we observe the same pattern of conductance and E_rev_ changes as in the presence of K^+^: the current increased continuously and the E_rev_ value exhibited only a transient positive excursion (Fig. 3B,D).

**Fig. 3.**
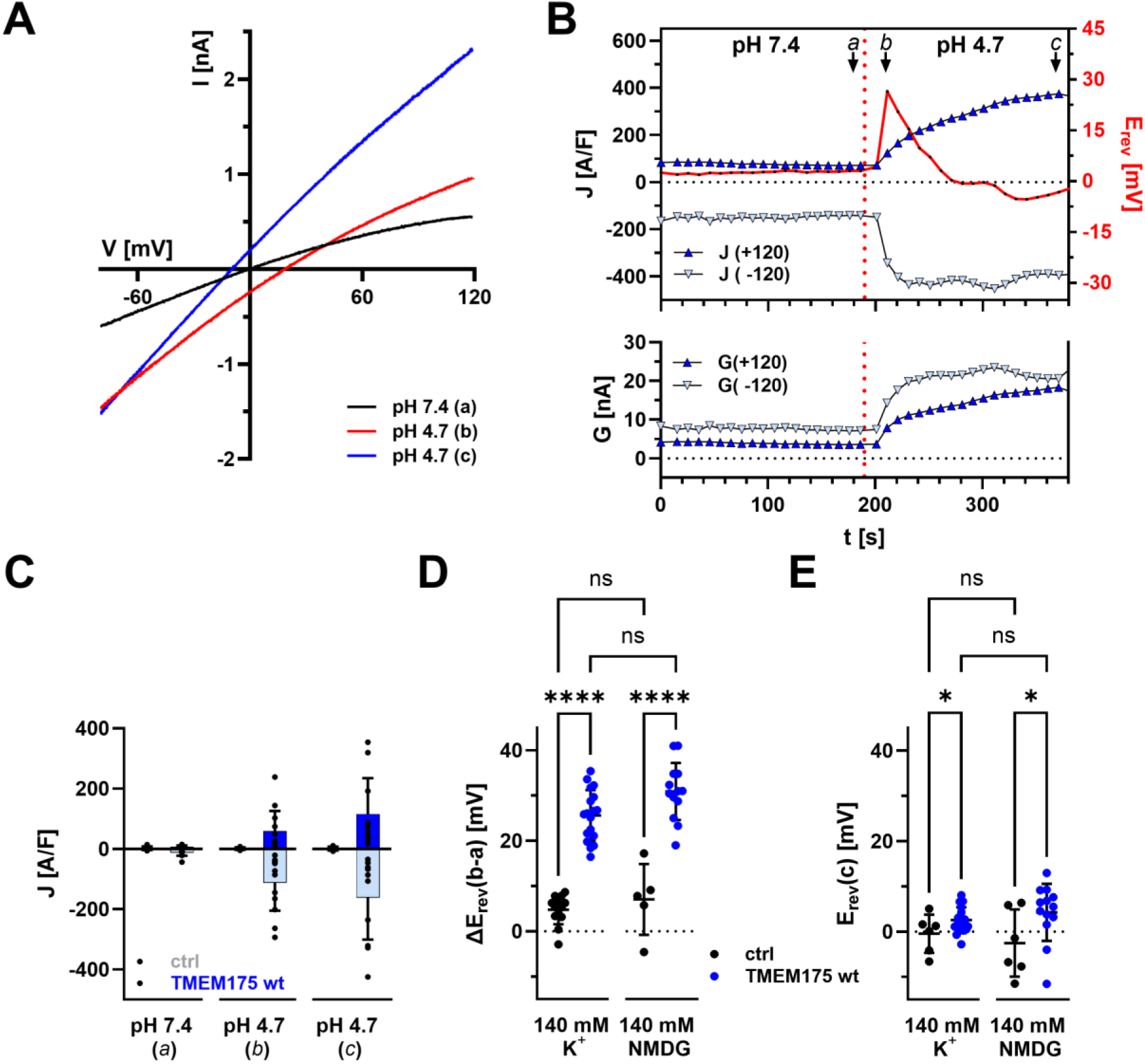
pH-induced conductance and transient positive E_rev_ excursion in TMEM175 expressing cells is K^+^ independent. **(A)** Representative current responses to voltage ramps from +120 mV to −120 mV of TMEM175 expressing HEK293 cells recorded shortly before (a, black), shortly after (b, red) and 3 min after (c, blue) a pH jump from 4.7 to 7.4 in symmetrical 140 mM NMDG-MS. The internal pH was 7.4. **(B)** Representative time-courses of reversal voltage (E_rev_) as well as current densities (upper graph) and chord conductance (lower graph) at +/- 120 mV of TMEM175 expressing HEK293 cells. Values were taken from voltage-ramp recordings as in (A). The external pH is indicated at the top. **(C)** Current densities at +/-120 mV of empty vector transfected (ctrl) and TMEM175 expressing cells at time points a, b and c in (B). Bars represent arithmetic mean ± SD. Values from individual recordings are shown as black closed circles. **(D)** Maximal change in reversal voltage E_rev_ in response to pH jump from 7.4 to 4.7 of empty vector transfected (black) and TMEM175 expressing (blue) HEK293 cells in symmetrical 140 mM K-MS (left) and 140 mM NMDG-MS (right). **(E)** Reversal voltages E_rev_ 3 min after pH jump (time point c in (B)) from 7.4 to 4.7 of empty vector transfected (black) and TMEM175 expressing (blue) HEK293 cells in symmetrical 140 mM K-MS (left) and 140 mM NMDG-MS (right). Data (D) and (E) show arithmetic means ± SD. Values from individual recordings are shown as closed circles. Statistical comparisons in (C-E) were made using analysis of variance (ANOVA) (ns: p > 0.05; *: p < 0.05, ****: p < 0.0001).

Collectively, the data show that the pH induced H^+^ currents are mechanistically independent of K^+^ currents. The decay of the E_rev_ value after an initial positive excursion is an inherent property of the H^+^ conductance. In this case the conductance properties are in the absence of K^+^ entirely dominated by H^+^ currents. In the presence of K^+^ the conductance is determined by a mixture of K^+^ and H^+^ conductance.

### TMEM175 is activated by acidification of the buffer on the luminal side

The experiments shown so far demonstrate that TMEM175 conducts protons and K^+^ after acidification of the extracellular solution. In the next experiments we address the question whether the increase in conductance after the pH jump simply reflects the change in the H^+^ driving force or whether it is the mixed result of pH-dependent gating plus an altered driving force. Therefore, we repeated experiments in the absence of K^+^ as in Fig. 3 with the difference that the pH of the pipette solution was set to 4.7. If the proton conductance of TMEM175 was not the result of proton-dependent activation from the luminal side, then TMEM175 expressing cells should show increased outward currents even at neutral extracellular pH. However, the results in Fig. 4 show that this is not the case. Instead, the magnitude of the inward and outward currents at an external pH of 7.4 is independent of the cytosolic pH value (Fig. 4A&C). Only acidification of the extracellular solution leads to a significant increase in TMEM175 conductance (Fig. 4A&B), which disappears again when the pH is changed back to 7.4 (Fig. 4B). Interestingly, the inward and outward currents at neutral external pH are significantly lower in the absence of K^+^ (i.e., in the presence of NMDG) than in the presence of K^+^ (Fig. 4C). This indicates that TMEM175 is able to conduct K^+^ even at neutral external pH.

**Fig. 4.**
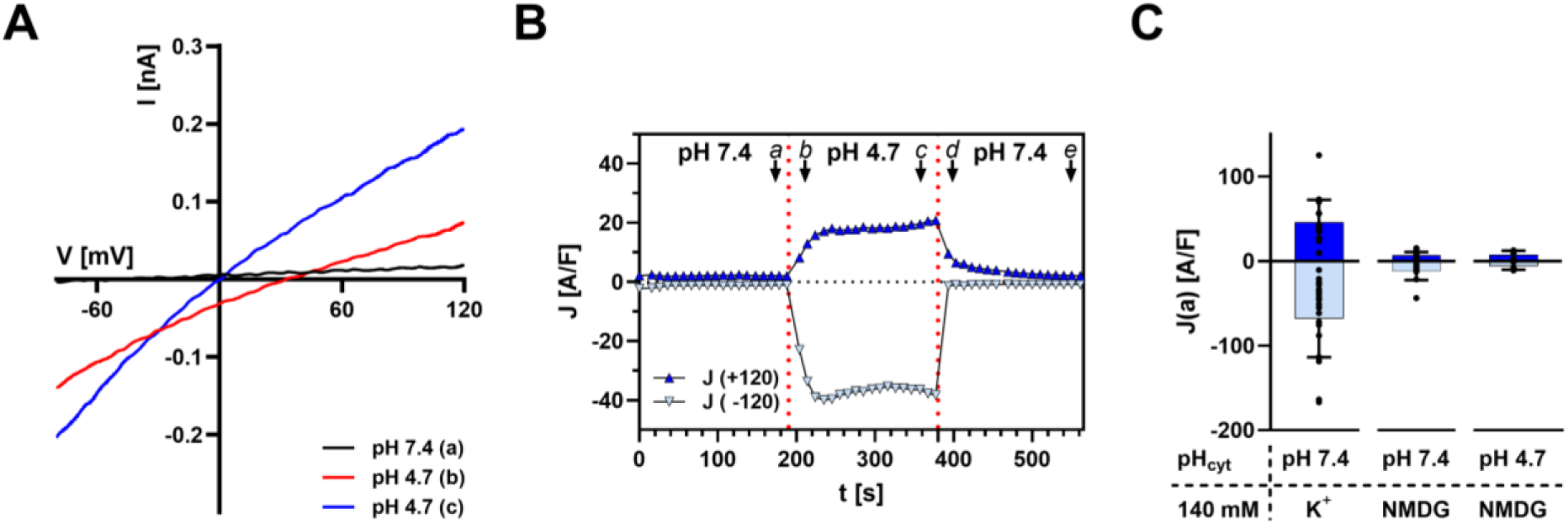
Acidification of the cytosolic side does not activate the H^+^ conductance of TMEM175. Whole-cell patch-clamp recordings of TMEM175 expressing HEK293 cells in symmetrical 140 mM NMDG-MS and an internal pH of 4.7. **(A)** Representative current responses to voltage ramps from +120 mV to −120 mV of a TMEM175 expressing HEK293 cell recorded shortly before (a, black), shortly after (b, red) and 3 min after (c, blue) an external pH jump from 4.7 to 7.4. **(B)** Representative time-course of current densities at +/-120 mV of TMEM175 expressing HEK293 cells. Values were taken from voltage-ramp recordings as in (A). The external pH is indicated at the top. **(C)** Current densities at +/-120 mV of TMEM175 expressing cells 3 min after the external pH jump from 7.4 to 4.7 in the presence of K-MS or NMDG-MS and a cytosolic pH of 7.4 or 4.7 as indicated. Bars represent arithmetic mean ± SD. Values from individual recordings are shown as black closed circles.

**Fig. 5.**
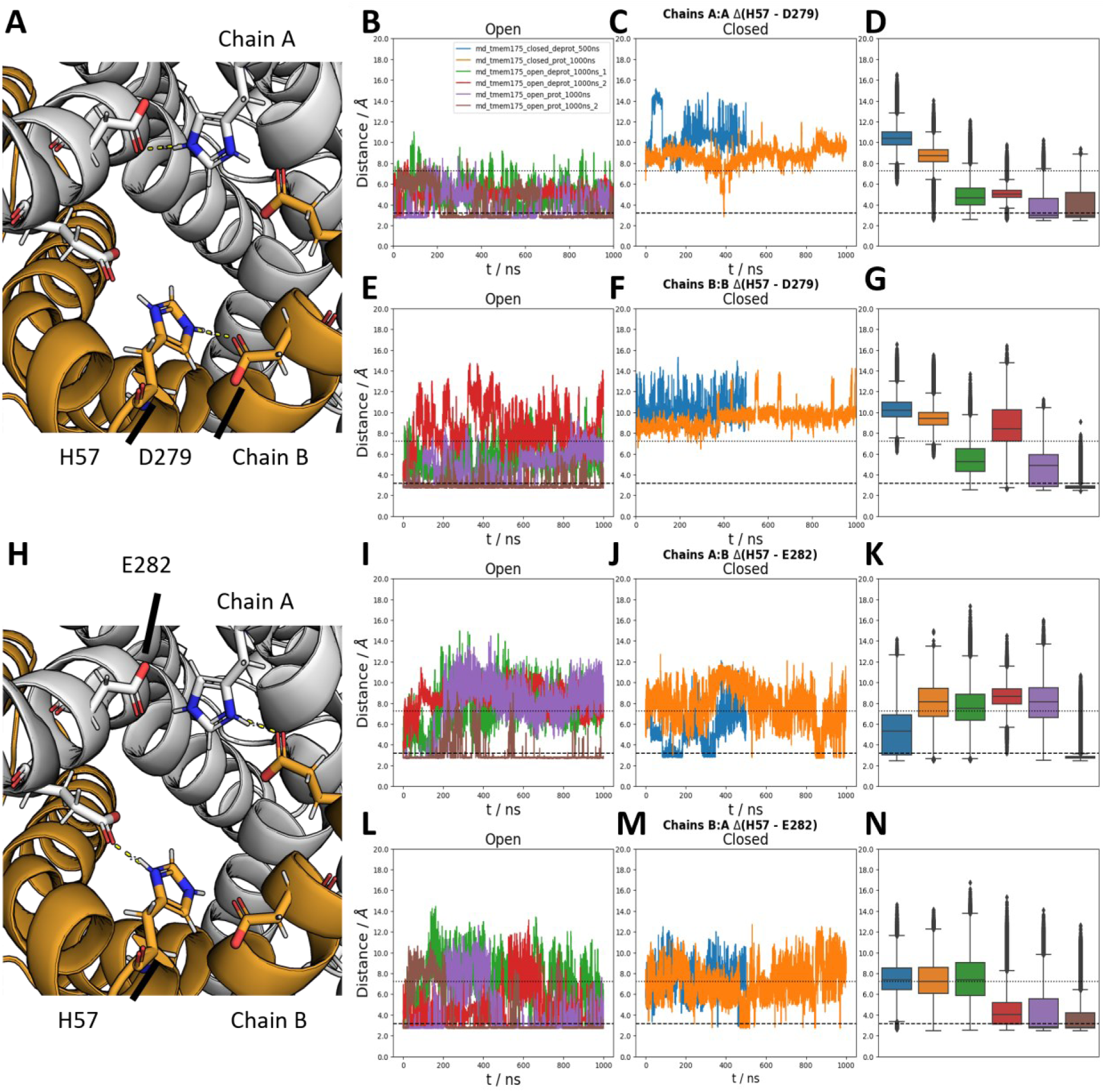
Salt bridge-compatible distances are observable between H57 and D279 for prolonged simulation time frames in simulations of the open-protonated TMEM175. Molecular images depict distances measured in trajectories for inter-monomeric H57-D279 distances (A) and inter-monomeric H57-E282 distances (H). The first two rows (B-G) show distances measured for H57-D279, while distances for H57-E282 are shown in the latter two rows (I-N). Row wise, data are separated according to distances measured between monomers, as detailed by the corresponding subtitles. Time series in the first column (B, E, I, L) depict the respective distances (B, E for H57-D279; I, L for H57-E282) measured for TMEM175 in the open state, with the following column depicting the corresponding distances drawn from simulations with the channel in the closed state (C, F, J, M). The moving average over 10 ns is depicted for visual clarity. Full, time-aggregated distance data for all simulations in the open/closed state for a given distance and monomer (pair) are shown as boxplot in the last column (D, G, K, N). Dotted lines denote the distance between H57 and its interaction partner measured in the cryoEM structure of TMEM175 in the open state, with dashed lines for the closed state. Lower distances for both simulation replicas with the channel in the open-protonated states (lilac, brown) are observable compared to the open-deprotonated simulations (green, red) and simulations in the closed-protonated/-deprotonated state (blue, orange).

Collectively, the results of these experiments suggest that the increase in current after acidification of the luminal side is determined by a combination of pH-sensitive gating and a change in driving force. The data further imply that the pH sensor is located on the luminal side of the channel, which is in HEK293 cells exposed to the extracellular solution.

### His57 plays a key role in sensing of the external pH

The data at this point predict that the TMEM175 channel has a pH-sensitive gate on the luminal side, which opens the channel in an acidic milieu. In search for a potential gating mechanism, we scanned the structure of the TMEM175 channel in the open and closed state (Oh et al., 2022) for potential candidate sites for such a gate on the luminal side. Under the assumption that this gate is operated by protonation/deprotonation at pH values around 7, we focused our attention on His57. In the open structure, this amino acid is in close proximity to two anionic amino acids namely D283 and E282 on the second monomer. We assumed that a protonated His57 can form salt bridge interactions with Glu282 and/or E279. This structural organization may guarantee an iris like structure with the effect that the entry to the channel pore is kept open. The hypothesis of a gating function is further supported by the finding that the critical His57 and the potential salt bridge partners are far apart in the closed channel structure. Hence, channel closing may be initiated by deprotonation of His57 with the consequent disruption of the stabilizing salt bridges and a closing of the gate.

To examine the plausibility of this hypothesis we performed molecular dynamics (MD) simulations with the open and the closed structure of TMEM175. In extended simulations, we monitored therefore the distance between the critical H57 in its protonated or de-protonated form with the adjacent intra-subunit glutamate D279 or with E282 on the second monomer. The simulations provide a complex picture showing that the amino acids in question can reach a critical distance for salt bridge formation in the open but not in the closed channel. In two simulations of the open channel, H57 remained stable for extended times at this critical distance to either E282 or D279 suggesting the formation of a stable salt bridge. The data also show that H57 achieves, in the open channel the critical distance to E282/D297 more frequently in its protonated than in its deprotonated form and exhibits also defined salt bridge formation with D297. Taken together the simulation results are compatible with the idea that H57 mediated salt bridge formation might operate in a pH dependent gate in the TMEM175 channel. The respective salt bridge could stabilize the open pore while its disruption as a consequence of the histidine deprotonation might destabilize this configuration. The variability of the simulation data and the finding that salt bridge formation in the open channel with protonated H57 is not obligatory and not always stable however suggests a more complex system which is not exclusively relying on a His57 mediated salt bridge formation. This assumption is in agreement with the finding that salt bridge formation occurs in the same simulation only between one of the two possible H57-E282 pairs. Hence other conformational changes in the protein are presumably contributing to this mechanism.

In the next step, we tested this hypothesis of a H57 mediated gating experimentally. The critical histidine (H57) was mutated to tyrosine. Tyrosine eliminates the titratable side chain of the histidine but maintains similar steric properties and retains the aromaticity in this position. TMEM175 and its H57Y mutant were expressed in HEK293 cells and currents recorded. The representative data as well as the corresponding analysis revealed that a pH jump from 7.4 to 4.7 elicits no increase in conductance in the mutant (Fig. 6A,B,F,G). Also, the typical positive excursion of the E_rev_ value recorded with the wild-type channel in response to the external pH jump was no longer observed in cells expressing the H57Y mutant (Fig. 6B,D,E). The results of these experiments are in good agreement with the hypothesis that His57 is part of a pH triggered gate in the TMEM175 channel. Thus, mutation of a single His abolishes the pH induced rise in conductance.

**Fig. 6.**
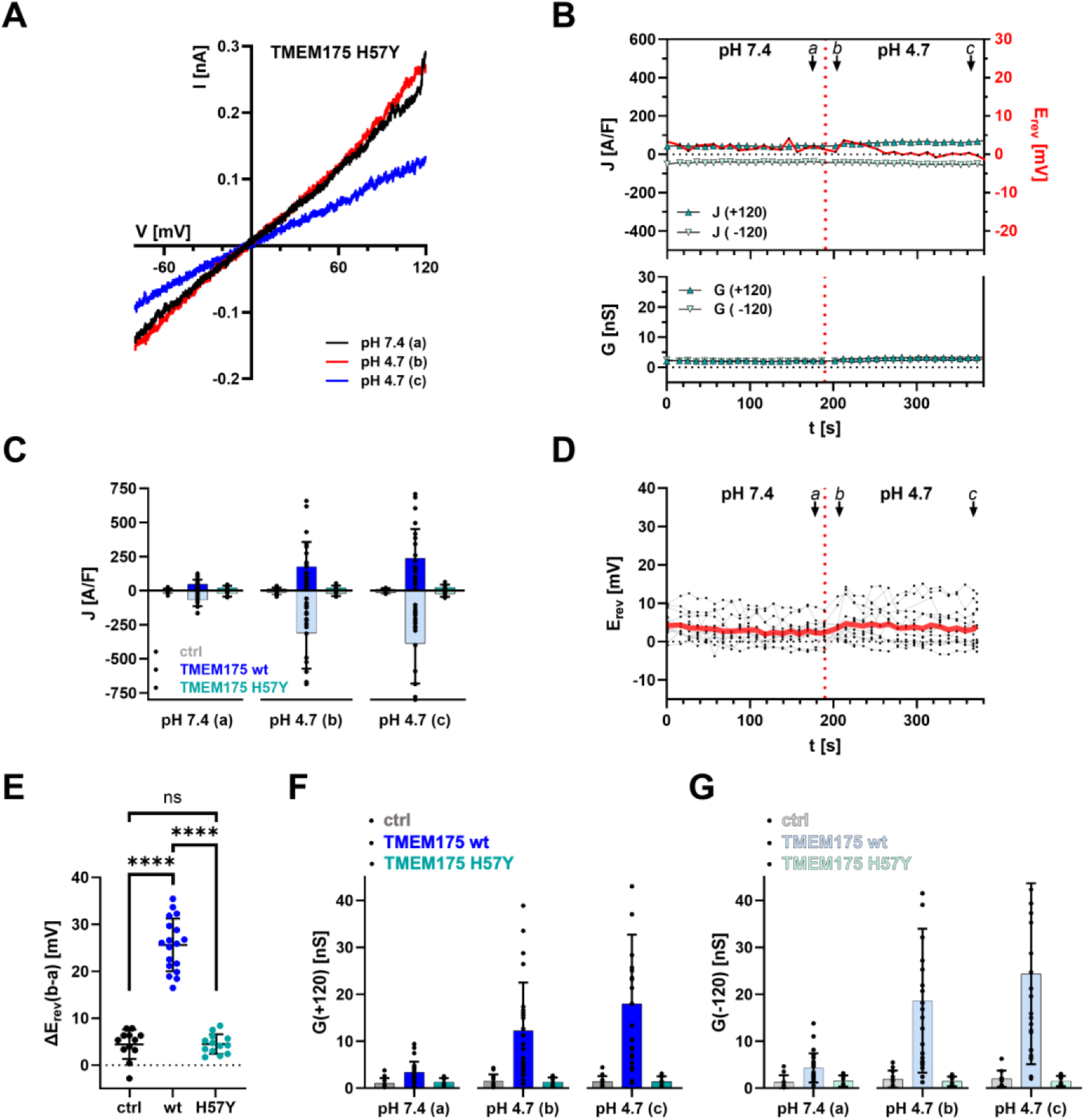
His57 is crucial for activation of TMEM175 by luminal acidification. **(A)** Representative current responses to voltage ramps from +120 mV to −120 mV of TMEM175 H57Y expressing HEK293 cells recorded shortly before (a, black), shortly after (b, red) and 3 min after (c, blue) a pH jump from 4.7 to 7.4 in the bath solution (luminal side). Bath and pipette solution contained 140 mM K-MS. The pH of the pipette solution was 7.4. **(B)** Representative time-courses of reversal voltage (E_rev_) as well as current densities (upper graph) and chord conductance (lower graph) at +/-120 mV of TMEM175 H57Y expressing HEK293 cells. Values were taken from voltage-ramp recordings as in (A). The external pH is indicated at the top. **(C)** Current densities at +/-120 mV at time points a, b and c in (B) for empty vector transfected, TMEM175 wt and TMEM175 H57Y expressing cells. Bars represent arithmetic mean ± SD. Values from individual recordings are shown as black closed circles. **(D)** Individual and averaged time-courses of E_rev_ of TMEM175 H57Y expressing HEK293 cells from pH jump experiments as in (B). **(E)** Maximal change in reversal voltage E_rev_ in response to pH jump from 7.4 to 4.7 of empty vector transfected (black), TMEM175 wt (blue) and TMEM175 H57Y (turquoise) expressing HEK293 cells. Data show arithmetic means ± SD. Values from individual recordings are shown as closed circles. Statistical comparisons were made using analysis of variance (ANOVA) (ns: p > 0.05; ****: p < 0.0001). **(F,G)** Chord conductance at +120 mV (F) and −120 mV (G) for empty vector transfected (gray), TMEM175 wt (blue) and TMEM175 H57Y (turquoise) expressing HEK293 cells at time points a, b and c in (B). Values were calculated from voltage-ramp recordings as in (A). Bars represent arithmetic mean ± SD.

Finally, we tested the relevance of our findings of a pH gated TMEM175 protein in its native environment of lysosomes. In control experiments we measured the membrane currents and the free running membrane voltage of isolated lysosomes in the “whole lysosome” configuration. The lysosomes were bathed in a buffer with pH 7.4 and internally dialyzed from the pipette with a buffer containing either a pH of 7.4 or 4.7. Fig. 7A shows a typical recording in which the current density at +120 mV and −120 mV was continuously monitored in two lysosomes after establishing the whole-lysosome configuration at time=0 by rupturing the membrane patch. In the representative recordings as well as in 5 similar experiments the respective currents were small and remained on this level over the 10 min of the recording irrespectively of the intra-lysosomal pH (Fig. 7A,C). When the same experiments were repeated in lysosomes overexpressing TMEM175 the respective inward and outward currents were significantly higher (Fig. 7A,C). When the lysosomal lumen was perfused with a neutral buffer the of inward and outward current densities reached, after ca. 3 min a stable value, which exceeded the control values by a factor of ca. 2 (Fig. 7C). In good agreement with the view of a pH gated activity of TMEM175 the respective current density settled gradually during dialysis with an acidic buffer at a stable value ca. 10 times higher than in controls (Fig. 7A,C). The difference in currents between the experimental conditions is also reflected in the free running membrane voltage V_mem_ of the organelles. While this value is irrespective of the pH around 5 mV in control cells it is smaller (2.7 ± 0.9 mV) or much higher (31.8 ± 4.0 mV) in lysosomes of TMEM175 expressing cells with neutral or acidic luminal pH, respectively (Fig. 7B).

**Fig. 7.**
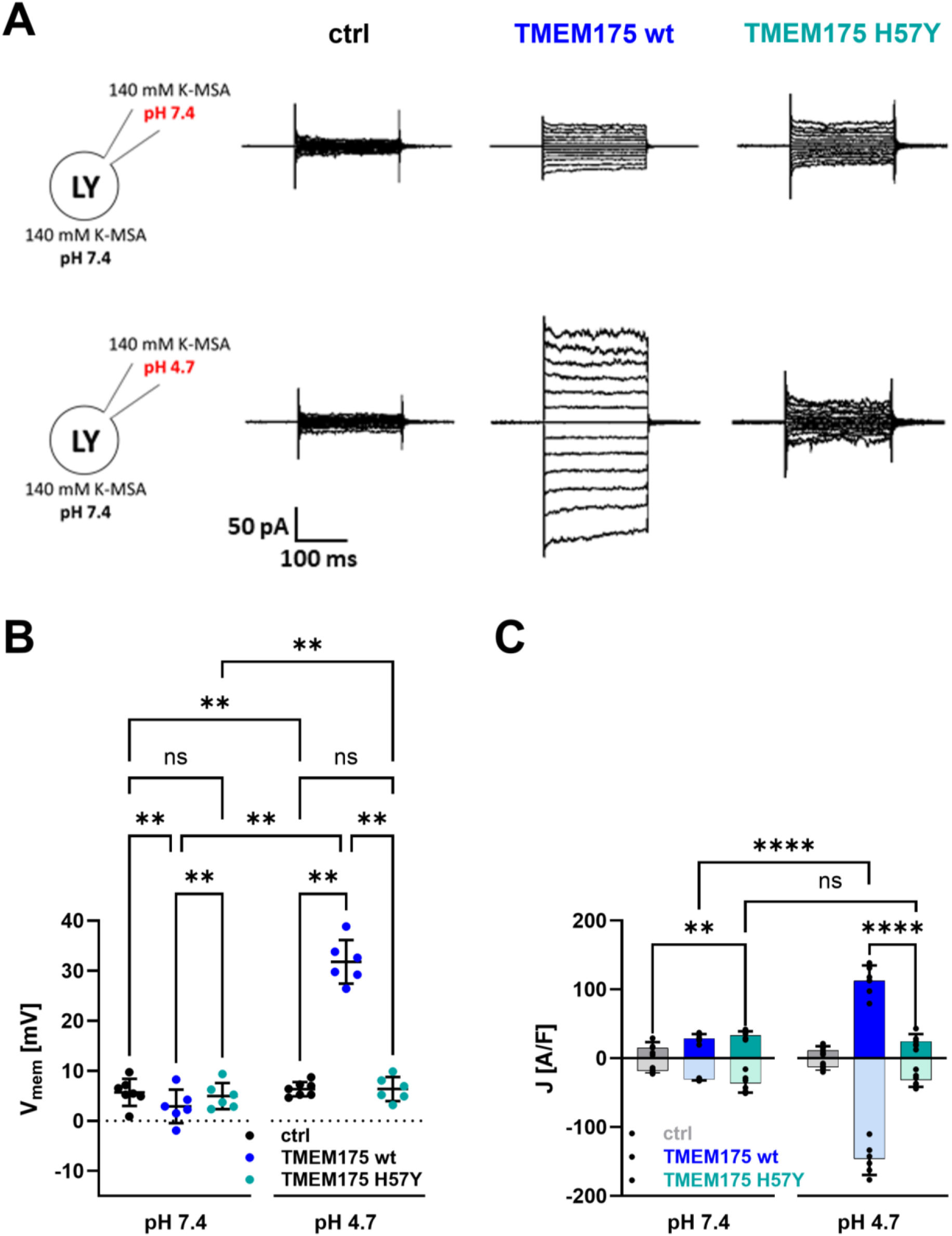
Endolysosomal patch-clamp recordings confirm the importance of His57 as luminal pH sensor. **(A)** Representative current responses to voltage step protocols from −120 mV to +120 mV in 20 mV increments of empty vector transfected (ctrl, left), TMEM175 wt (middle) and TMEM175 H57Y expressing HEK293 cells (right) recorded 3 min after achievement of whole-endolysosome configuration. The lysosomes were bathed in a buffer with pH 7.4 and internally dialyzed from the pipette with a buffer containing either a pH of 7.4 (upper row) or 4.7 (lower row). **(B)** Free running membrane potential of lysosomes from empty vector transfected cells (ctrl, black), hTMEM175 wt (blue) and TMEM175 H57Y expressing cells with internal pH 7.4 or pH 4.7 expressing cells measured the same way as in (A) recorded 3 min after achievement of whole-endolysosome configuration. Data show arithmetic means ± SD. Values from individual recordings are shown as closed circles. **(C)** Current densities at +/-120 mV of empty vector transfected (ctrl, grey) and TMEM175 expressing cells. Bars represent arithmetic mean ± SD. Values from individual recordings are shown as black closed circles. Statistical comparisons in (B) and (C) were made using analysis of variance (ANOVA) (ns: p > 0.05; *: p < 0.05, ****: p < 0.0001).

Important to mention here is that the steady state E_rev_ is also in TMEM175 over-expressing lysosomes with an acidic lumen well below the theoretically expected Nernst voltage for H^+^. These findings confirm that the channels exhibit in lysosomes similar functional features to those in the plasma membrane of HEK293 cells. To test the relevance of the H57 residue for pH-sensitive gating the mutant was expressed in lysosomes and the mutant currents monitored. The representative recordings from single lysosomes and the corresponding mean values from multiple repetitions of these experiments show that the mutation completely abolishes the positive effect of acidic pH on channel conductance (Fig. 7A,C). The current densities and the values of the free running membrane voltage in experiments with lysosomes expressing the mutant and perfused with neutral or acidic luminal pH, respectively, are indistinguishable from control recordings (Fig. 7B). These results confirm the data from complementary recordings of the mutant in the plasma mebrane of HEK293 cells and underpin the relevance of the luminal H57 residue for TMEM175 gating. A more fine-grained comparison of the current density and membrane voltage data furthermore shows that the mutant generates current densities which are significantly higher than the control values measured in lysosomes not overexpressing the TMEM175 channel (Fig. 7A,C). This suggests that the channel is not fully closed and may still conduct some K^+^ and/or H^+^ ions.

### Conclusions

The systematic examination of pH induced conductance and voltage changes in TMEM175 expressing cells and lysosomes argue for a pH sensitive gate on the luminal side of the channel. The latter involves a critical Histidine and its salt bridge interactions with adjacent anionic amino acids for stabilizing the channel open conformation as a function of the protonation state of this histidine. Upon activation, the TMEM175 channel conducts, in a K^+^ independent manner preferentially H^+^. These proton currents generate a rapid transmembrane acidification and thus a progressive decrease in the driving force for protons. The latter occurs with such a high velocity that TMEM175 mediated H^+^ currents exhibit equilibrium voltages which are already within seconds well below the theoretically expected Nernst potential for H^+^. The pH dependent gating, the defined selectivity for H^+^ over K^+^ as well as the kinetics of H^+^ currents with their negative feedback on the driving force, provide missing information for a complete understanding of the spatial-temporal functions of this protein in lysosomes.

## Materials and Methods

### Cloning

The expression vector pEGFP-N2 was double restriction digested with NotI and XhoI, removing the EGFP gene. The sequence coding for human TMEM175 was then amplified from pcDNA3.1-hTMEM175_STOPSTOP (Brunner et al., 2020) using primer pair 1 (table 1) and the amplicon was inserted downstream of the CMV promoter sequence via Gibson assembly with NEBuilder® HiFi DNA Assembly Master Mix (NEB, Ipswich, MA, USA) into the pEGFP-N2 backbone. Reaction products were used for heat-shock transformation of chemically competent DH5α *E. coli*. Selection of positive clones was performed on LB Agar plates containing 25 µg/ml kanamycin. In addition, site-directed mutagenesis (SDM) was performed on this vector using primer pair 2 (table 1) to induce the point mutation H57Y into TMEM175. Success of cloning and SDM was confirmed by sequencing.

**Table 1:**
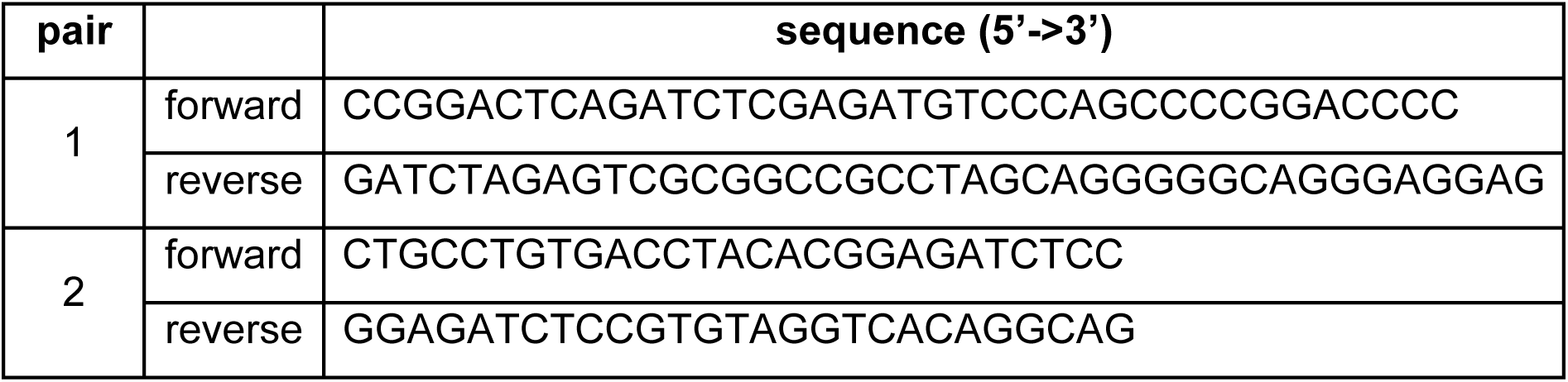
Primers used for cloning and SDM.

### Cell culture

For functional expression of TMEM175 and TMEM175 mutants HEK293 cells were transfected 16 to 24 hours before the start of patch-clamp experiments using TransfeX™ Transfection Reagent (LGC Standards GmbH, Wesel, Germany) according to the manufacturer’s instructions. HEK293 cells were grown at 37°C in a humidified 95% air/5% CO_2_ incubator in Dulbecco’s Modified Eagle Medium (DMEM; Gibco) supplemented with 10% v/v heat-inactivated fetal bovine serum, 100 U/ml penicillin G, 100 μg/ml streptomycin sulfate and 2 mM L-glutamine (all from Invitrogen). After reaching approximately 80% confluence mammalian cells were (co-)transfected in a 35 mm petri dish with 1 µg of the plasmid carrying the gene of interest and (if necessary) 1 µg of empty pEGFP-N2 vector to be able to identify transfected cells via eGFP fluorescence.

### Plasma membrane patch-clamp electrophysiology

On the day of the experiment, transfected HEK293 were separated by trypsinization, seeded at low density on 10 mm coverslips, and then incubated for 2 to 4 hours to allow adhering of cells on the glass surface. For patch-clamp experiments, coverslips were then transferred to a perfusion chamber filled with bath solution and placed on the stage of an inverted microscope. Transfected cells were identified by the fluorescence of co-expressed eGFP. Patch-clamp experiments were performed at room temperature (20-25°C) with an EPC-9 amplifier (HEKA Elektronik, Lambrecht, Germany) in the whole-cell configuration. Patch-pipettes were pulled from borosilicate capillaries (DWK Life Sciences, Milville, NJ, USA) using a single stage glass microelectrode puller (PP-830, Narishige Group, Tokyo, Japan) resulting in 2-3 Mohm resistances. Capillaries were coated with Sigmacote® (Merck KgaA, Darmstadt, Germany) and baked after pulling at 65°C for 45 min. The bath electrode was protected with a 1.5% agar bridge. The agar bridges were prepared from a solution of 200 mM KCl. Currents were recorded with a 10 kHz low-pass Bessel filter and sampled with a frequency of 20 kHz.

To determine the time-dependent activation of TMEM175 after pH activation a symmetrical ramp protocol was used. A holding potential of +120 mV is held for 25 ms followed by a voltage ramp for 150 ms to a holding potential of −120 mV for another 25 ms. Afterwards the voltage was continuously increased to +120 mV over 150 ms.

Free running Membrane potentials were measured in current-clamp mode at zero current. For current-clamp measurements, the ‘Gentle CC-Switch’ option in PatchMaster was deactivated, so that the holding potential is not retained when switching from voltage-clamp to current-clamp, but changes directly to 0 pA. Liquid junction potentials (LJPs) were calculated using JPCalcWin (UNSW, Sydney, Australia) and subtracted post recording. Data were stored with PatchMaster (HEKA Elektronik, Lambrecht, Germany) and analyzed with FitMaster (HEKA Elektronik, Lambrecht, Germany).

Unless otherwise stated, standard pipette solution contained in mM: 140 potassium methanesulfonate (KCH_3_SO_3_; K-MS), 10 TEA-OH, 5 cesium methanesulfonate (CsCH_3_SO_3_; Cs-MS), 2.5 MgCl_2_, 5 4-(2-hydroxyethyl)-1-piperazineethanesulfonic acid (HEPES), 1 ethylene glycol-bis(β-aminoethyl ether)-N,N,N′,N′-tetraacetic acid (EGTA). pH and osmolarity were adjusted to 7.4 with 1 M KOH and 300 mOsmol/kg with D-mannitol, respectively.

The standard bath solution at pH 7.4 contained unless stated otherwise in mM: 140 K-MS, 10 TEA-OH, 5 Cs-MS, 0.5 MgCl_2_, 1.8 CaCl_2_, 5 HEPES. pH and osmolarity were adjusted to 7.4 with 1 M KOH and 290 mOsmol/kg with D-mannitol, respectively. The standard bath solution at pH 4.7 contained unless stated otherwise in mM: 130 K-MS, 10 TEA-OH, 5 Cs-MS, 0.5 MgCl2, 1.8 CaCl2, 10 potassium acetate (KOAc) and 10 acetic acid (HOAc). pH and osmolarity were adjusted to 4.7 with 1 M KOH or metahesulfonic acid (MSA) and 300 mOsmol/kg with D-mannitol, respectively.

### Endolysosomal patch-clamp electrophysiology

Manual whole-endolysosomal electrophysiology was performed in isolated enlarged endosome/endolysosome vacuoles using a modified patch-clamp method as described previously (Chen et al., 2017). HEK293 cells were plated onto poly-L-lysine (Sigma)-coated glass coverslips, grown over night and transiently transfected for 16–24 hours with plasmids using TurboFect (ThermoFisher) according to the manufacturer’s instructions. As a next step, cells were treated for >8 hrs with 1 µM apilimod, a lipophilic polycyclic triazine that can selectively increase the size of endosomes and endolysosomes. Large vacuoles/endolysosomes were observed in most apilimod-treated cells. Transfected cells were identified by the fluorescence of co-expressed eGFP. Glass pipettes for recording were polished and had a resistance of 4–8 MΩ. Patch-clamp experiments were performed at room temperature (20-25°C) with an EPC-10 amplifier (HEKA Elektronik, Lambrecht, Germany) and PatchMaster acquisition software (HEKA) in the whole-endolysosome configuration on manually isolated enlarged vacuoles. In brief, a patch pipette was pressed against a cell and quickly pulled away to slice the cell membrane. This allowed enlarged endolysosomes to be released into the recording chamber and identified by monitoring EGFP fluorescence. After formation of a gigaseal between the patch pipette and an enlarged endolysosome, capacitance transients were compensated. Voltage steps of several hundred millivolts with millisecond duration(s) were then applied to break the patched membrane and establish the whole-endolysosome configuration. Data were collected using the same patch solutions as already described for the plasma membrane patch clamp experiments. To measure steady-state currents a step protocol was used. After holding 0 mV for 100 ms, test pulses starting from −120 mV were applied with a voltage increment of 20 mV for 200 ms. A holding potential of 0 mV was then clamped for further 100 ms.

### Data analysis

Data was exported using PatchMaster and FitMaster (HEKA Elektronik, Lambrecht, Germany) and sorted as well as analyzed in Excel (Microsoft; Redmond, Washington, USA) and MATLAB (MathWorks; Natick, Massachusetts, USA). Statistical analyses were carried out in GraphPad Prism.

Data are presented as mean ± standard deviation (SD) from at least three independent experiments. Statistical comparisons were made using analysis of variance (ANOVA) and Student’s t test. P values of < 0.05 were considered statistically significant. *: p < 0.05; **: p < 0.01; ***:p < 0.001; ****: p < 0.0001.

### Structure preparation and modelling

hTMEM175 cryoEM structures in the open (PDB accession number: 7UNL) and closed state (PDB accession number: 7UNM) were used as basis for modeling (Oh et al., 2022). To fill the existing gaps between residue 173 and residue 254 in both structures, the Alphafold2-multimer model as implemented by ColabFold was used to generate five individual structures each (Evans et al., 2021; Mirdita et al., 2022). Those were energy-minimized according to the standard procedure in ColabFold (Mirdita et al., 2022).

The two structures with the lowest pLDDT and PAE scores were prepared for MD simulations. The alpha-helical parts of the structure show a pLDDT of about 70 while some unstructured parts are predicted with a pLDDT score below 70. Scores above 70 are commonly deemed acceptable, while lower pLDDTs are also associated with unstructured stretches in other studies (Tunyasuvunakool et al., 2021; Wilson et al., 2022). As the main region of interest is located proximate to H57, a lower fidelity is acceptable in modeling.

Using CHARMM-GUI, the open and closed structures were embedded in POPC lipid bilayers, for a final KCl concentration of 150 mM (J. Lee et al., 2016).

Both the open and closed structure were prepared with H57 in the protonated and deprotonated form.

### Equilibration and MD simulations

The Amber99sb*-ILDN force field was used in combination with Cheatham-Joung parameters for Na^+^/K^+^ and Cl^-^ and the TIP3P water model (Best & Hummer, 2009; Jorgensen et al., 1983; Joung & Cheatham, 2008; Lindorff-Larsen et al., 2010). To allow for a timestep of 4 fs in all simulations, the partially coarse-grained Berger-derived lipid parameters were used to model POPC, together with a LINCS-constraint on all bonds and virtual sites for hydrogen atoms (Berger et al., 1997; Hess et al., 1997).

This combination was validated for higher simulation timesteps and was successfully employed for a timestep of 4 fs in several publications (Feenstra et al., 1999; Saponaro et al., 2021). Electrostatic interactions were treated with the particle-mesh Ewald method with a 1 nm real space cutoff. Van-der-Waals interactions were cut off above 1 nm. In all simulations, the temperature was held constant at 310 K with the V-rescale thermostat (Bussi et al., 2007). A pressure of 1 bar was maintained using the Berendsen (equilibration) and Parinello-Rahman (production simulations) barostats (Berendsen et al., 1984; Parrinello & Rahman, 1981).

All simulations were run with Gromacs 2021.5 (Abraham et al., 2015). Prior to production runs, the following energy minimization/equilibration scheme was employed: Energy minimization was conducted for 2000 steps with the steepest-descent integrator and a step width of 0.1 Å, followed by 100 ps in the NVT ensemble with restraints for protein sidechains and backbone residues of F_c_ = 1,000 kJ/mol acting on sidechains and backbone atoms, respectively. This was followed by subsequent NPT simulations with gradually lifted restraints on backbon/sidechains, as previously described in Saponaro et al 2021(Saponaro et al., 2021).

Production simulations were run over 0.5 µs-1 µs. For the most relevant states (open-prot and open-deprot), three separate simulations over 1µs were run.

### Analysis of MD simulations

Obtained MD trajectories were analyzed using biotite (Version 0.41.0) together with the Gromacs (v. 2021.5) analysis tools (Abraham et al., 2015; Kunzmann et al., 2023). Various other Python libraries were used for general data analysis and visualization; a complete list of all used software and libraries is shown in the Supplement.

## Acknowledgements

This work was supported by DFG (Forschungsimpuls CytoTransport, project ID 528562393 – FIP26) and grants HA5261/6-1 to K. Hamacher, TH558/34-1 to G. Thiel, and DFG4315/2-2, DFG/ANR4315/6-1, DFG4315/7-1, SFB1328 A21, SFB/TRR152 P04 and GRK2338 P08 to C. Grimm.

