## Supplementary Figures S1-S3 for "Proton selective conductance and gating of lysosomal cation channel TMEM175"

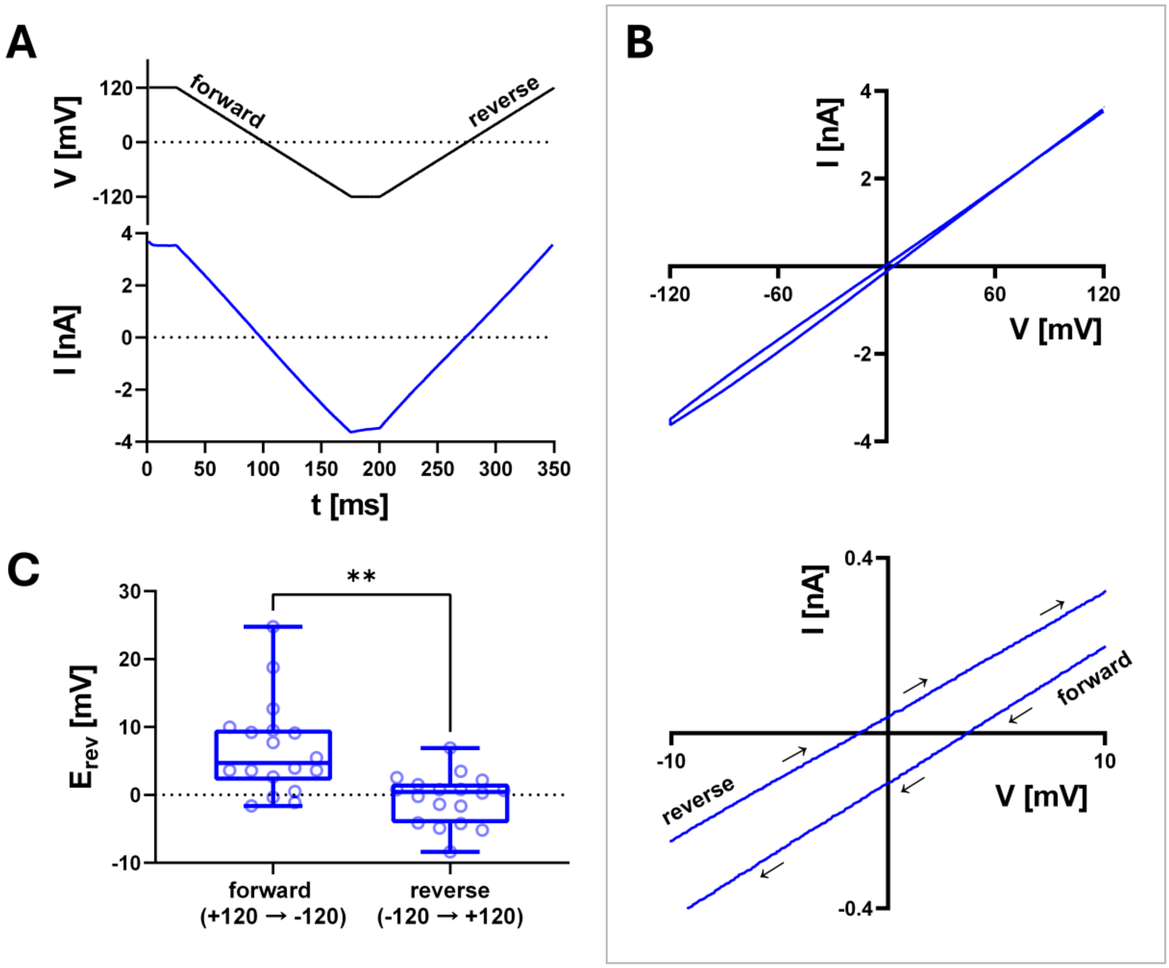


**Fig.S1 Voltage ramp evoked currents cause shifts in current reversal potential. (A)** Voltage-ramp protocol (top) and representative current response (bottom) of a TMEM175 expressing cell 3 min after acidification of the external buffer. Experiment was performed as in Fig. 1. **(B)** Top: full current response from (A) plotted against the applied voltage. Bottom: Magnification of currents to illustrate the shift in the reversal potential $E_{rev}$ between forward and reverse ramp. The direction in which the voltage was changed during the forward and reverse ramp is indicated by arrows. **(C)** Reversal potentials of the forward and reverse ramps from current responses as in (A). Statistical comparison was made using a paired two-tailed Student's t-test (**: $p < 0.01$).

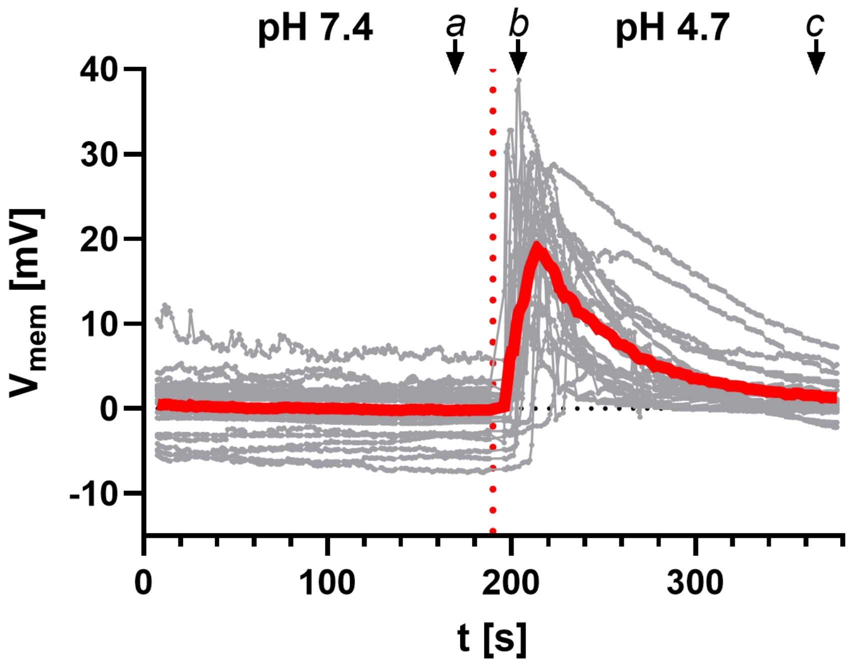


**Fig.S2 Acidification of the bath solution induces transient shift in free running membrane voltage to positive values in TMEM175 expressing HEK293 cells.** Individual (gray) and averaged (red) time-courses of the free running membrane voltage ($V_{mem}$) of TMEM175 expressing HEK293 cells with symmetrical 140 mM K-MS. The internal pH was 7.4. The external pH is indicated at the top. Recordings were performed in the current-clamp mode.

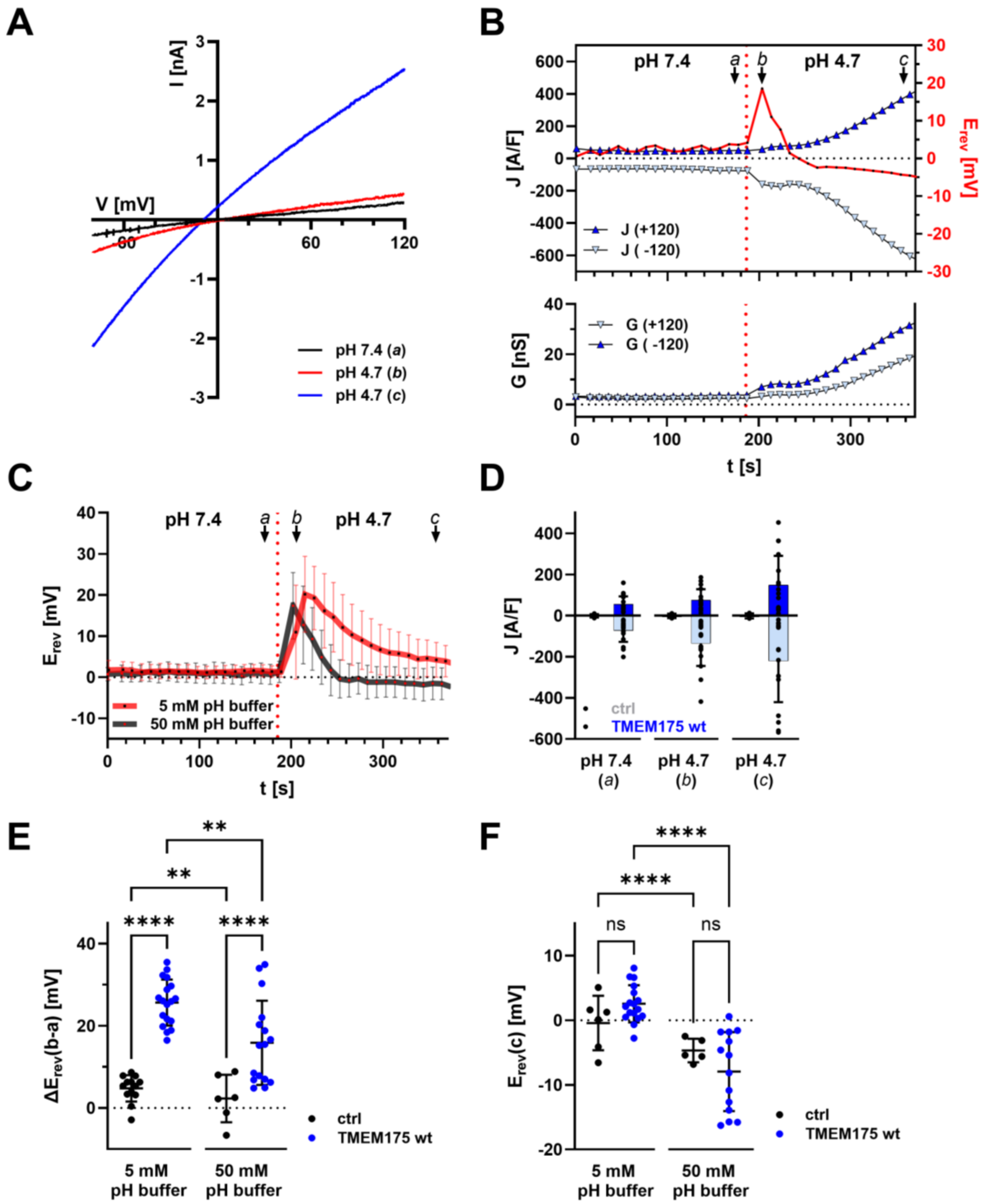


**Fig.S3 Increase of the pH-buffer concentration by a factor of 10 accelerates the backshift of $E_{rev}$ towards 0 mV after acidification of the external solution.**

**(A)** Representative current responses to voltage ramps from +120 mV to -120 mV of TMEM175 expressing HEK293 cells recorded shortly before (a, black), shortly after (b, red) and 3 min after (c, blue) a pH jump from 4.7 to 7.4 in the bath solution (luminal side). Bath and pipette solution contained 50 mM pH buffer (HEPES for pH 7.4, KOAc/HOAc for pH 4.7). The pH of the pipette solution was 7.4. **(B)** Representative time-courses of reversal voltage ($E_{rev}$) as well as current densities (upper graph) and chord conductance (lower graph) at +/-120 mV of TMEM175 expressing HEK293 cells in the presence of 50 mM of the respective pH buffer. Values were taken from voltage-ramp recordings as in (A). The external pH is indicated at the top. **(C)** Mean time-courses of $E_{rev}$ of TMEM175 expressing HEK293 cells from pH jump experiments as in (B) with 5 mM (red) or 50 mM (gray) of the respective pH buffer. Error bars represent SD. **(D)** Current densities at +/-120 mV at time points a, b and c in (B) for empty vector transfected and TMEM175 expressing cells in the presence of 50 mM of the respective pH buffer. Bars represent arithmetic mean ± SD. Values from individual recordings are shown as black closed circles. **(E)** Maximal change in reversal voltage $E_{rev}$ in response to pH jump from 7.4 to 4.7 of empty vector transfected (black) and TMEM175 expressing cells (blue) with 5 mM (left) and 50 mM (right) of the respective pH buffer. **(F)** Reversal voltage $E_{rev}$ 3 min after the pH jump (time point c in (B)) of empty vector transfected (black) and TMEM175 expressing cells (blue) with 5 mM (left) and 50 mM (right) of the respective pH buffer. Data in (E) and (F) show arithmetic means ± SD. Values from individual recordings are shown as closed circles. Statistical comparisons were made using analysis of variance (ANOVA) (ns: $p > 0.05$; **: $p < 0.01$;****: $p < 0.0001$).
